# Identification of receptor-binding domains of Bacteroidales antibacterial pore-forming toxins

**DOI:** 10.1101/2025.01.06.631476

**Authors:** Sofia Borgini, Bogdan Iorga, Didier Vertommen, Jean-François Collet, Frédéric Lauber

**Affiliations:** de Duve Institute, Université Catholique de Louvain, Avenue Hippocrate 75, 1200 Brussels, Belgium; WELBIO, Avenue Hippocrate 75, 1200 Brussels, Belgium; Université Paris-Saclay, CNRS UPR 2301, Institut de Chimie des Substances Naturelles, 91198 Gif-sur-Yvette, France

**Keywords:** Gram-negative bacteria, Bacteroidales, BSAP, pore-forming toxin, MACPF, protein-protein interaction, receptor, outer membrane, competition

## Abstract

Bacteroidales are abundant Gram-negative bacteria present in the gut microbiota of most animals, including humans, where they carry out vital functions for host health. To thrive in this competitive environment, Bacteroidales use sophisticated weapons to outmatch competitors. Among these, BSAPs (Bacteroidales Secreted Antimicrobial Proteins) represent a novel class of bactericidal pore-forming toxins that are highly specific to their receptor, typically targeting only a single membrane protein or lipopolysaccharide. The molecular determinants conferring this high selectivity remain unknown. In this study, we therefore investigated the model protein BSAP-1 and determined which of its domains is involved in providing receptor specificity. We clearly demonstrate that receptor recognition is entirely driven by the C-terminal domain (CTD) of BSAP-1 using a combination of *in vivo* competition assays and *in vitro* protein binding studies. Specifically, we show that deletion of the CTD abrogates BSAP-1 bactericidal activity by preventing receptor binding, while grafting the CTD to unrelated carrier proteins enables CTD-driven interaction with the BSAP-1 receptor. Building upon this discovery, we show that BSAPs can be categorized according to the structure of their CTD and that BSAPs within the same cluster are likely to target the same type of receptor. Additionally, we show that the CTD of BSAP-1 can be repurposed to generate probes for fluorescent labelling of membrane proteins in live cells. In summary, our research demonstrates that BSAP receptor recognition is driven by their CTD and that these can be engineered to develop novel tools for the investigation of Bacteroidales biology.

## Introduction

Bacteroidales are efficient long-term colonisers of the human intestine and are, on average, the most abundant Gram-negative bacteria in a healthy gut microbiota (1–3). They play pivotal roles in human health thanks to their positive anti-inflammatory (4,5), immunomodulatory (6) and metabolic properties (7,8). To secure their ecological niche in this highly competitive environment, Bacteroidales employ numerous mechanisms to directly antagonize opponents. These include direct injection of toxins via the cell contact-dependent Type 6 Secretion System (T6SS) (9–11) as well as production of antibacterial molecules such as BSAPs (Bacteroidales Secreted Antimicrobial Proteins) (12–15), bacteroidetocins (16,17), and the Fab1 (18), BfUbb (19,20) and BcpT (21) toxins. Among these, the recently identified BSAPs are widespread throughout Bacteroidales and have been shown to actively drive microbiota strain selection, structuring the intestinal flora in space and time (13,14,22,23). Deciphering their mechanism of action at a molecular level therefore holds promise to better understand how a healthy human gut microbiota is established and maintained over time.

BSAPs are surface-exposed lipoproteins, *e.g.* globular proteins anchored to the outside of the cell by an invariable lipidated N-terminal cysteine (24) and targeted to the cell surface by a specific lipoprotein export signal (LES) downstream of that residue (25,26). In addition, all BSAPs possess a conserved Membrane Attack Complex/Perforin (MACPF) domain (Pfam PF01823) (27) and belong to the wider MACPF/CDC superfamily of proteins (28) that includes cholesterol-dependent cytolysins (CDCs) (29) produced by Gram-positive pathogens and eukaryotic MACPF proteins involved in innate immunity (30,31). BSAPs therefore represent a new class of bacterial pore-forming toxins (PFTs) and are predicted to kill their target cells by formation of large lytic membrane pores (32). Indeed, investigation of BSAP-1 (BF638R_1646, GenBank: CBW22174.1), expressed by *Bacteroides fragilis* 638R and targeting a specific outer membrane β-barrel receptor protein (hereafter B1R^S^) present in a subset of *B. fragilis* strains, revealed that exposure of BSAP-1-sensitive cells to purified BSAP-1 led to a rapid uptake of the cell-impermeable DNA dye propidium iodide (12). This strongly suggests that BSAP-1 disrupts membrane integrity of sensitive cells and that BSAP-1 bactericidal activity results from lytic pore formation, similar to other MACPF/CDC proteins. By analogy with characterized members of this group of proteins, the conserved MACPF domains is the likely driving force for BSAP subunit oligomerization and pore formation (28). The precise architecture and assembly process of a BSAP lytic pore remain however to be determined.

A striking property of BSAPs is that, unlike other members of the MACPF/CDC superfamily, BSAPs are highly specific to their receptor and typically only target a single outer membrane β-barrel protein (12,15) or lipopolysaccharide (LPS) O-antigen (13,14). Due to this high specificity, BSAPs have so far only been described in the context of intra-species competition, making them the first example of bacterial MACPF proteins possessing bactericidal activity. This contrasts with other MACPF and CDC proteins that generally display inter-kingdom activity and use widespread receptor types, such as cholesterol in the case of CDCs (27). The molecular determinants conferring BSAPs this uniquely high selectivity remain currently unknown.

To address this gap of knowledge, we investigated BSAP receptor recognition and binding. Following maturation and surface translocation, BSAPs are typically composed of three domains: a disordered N-terminal region harboring the lipidated cysteine and LES export sequence; the highly conserved MACPF domain; and one to several C-terminal domains (CTD) of unknown function (Fig. 1A) (32). Because the MACPF domain is predicted to be essential for BSAP subunit oligomerization and lytic pore formation (28), is it unlikely to additionally be involved in providing specificity to receptors as diverse as proteins or glycans. Instead, sequence conservation analyses showed that CTDs vary greatly, resulting in a wide array of different BSAP architectures (14,32). As BSAP receptors are likewise highly variable, we therefore hypothesized that high CTD diversity directly correlates with that of BSAP receptors, and that CTDs dictate receptor specificity and are essential for receptor binding.

**Figure 1.**
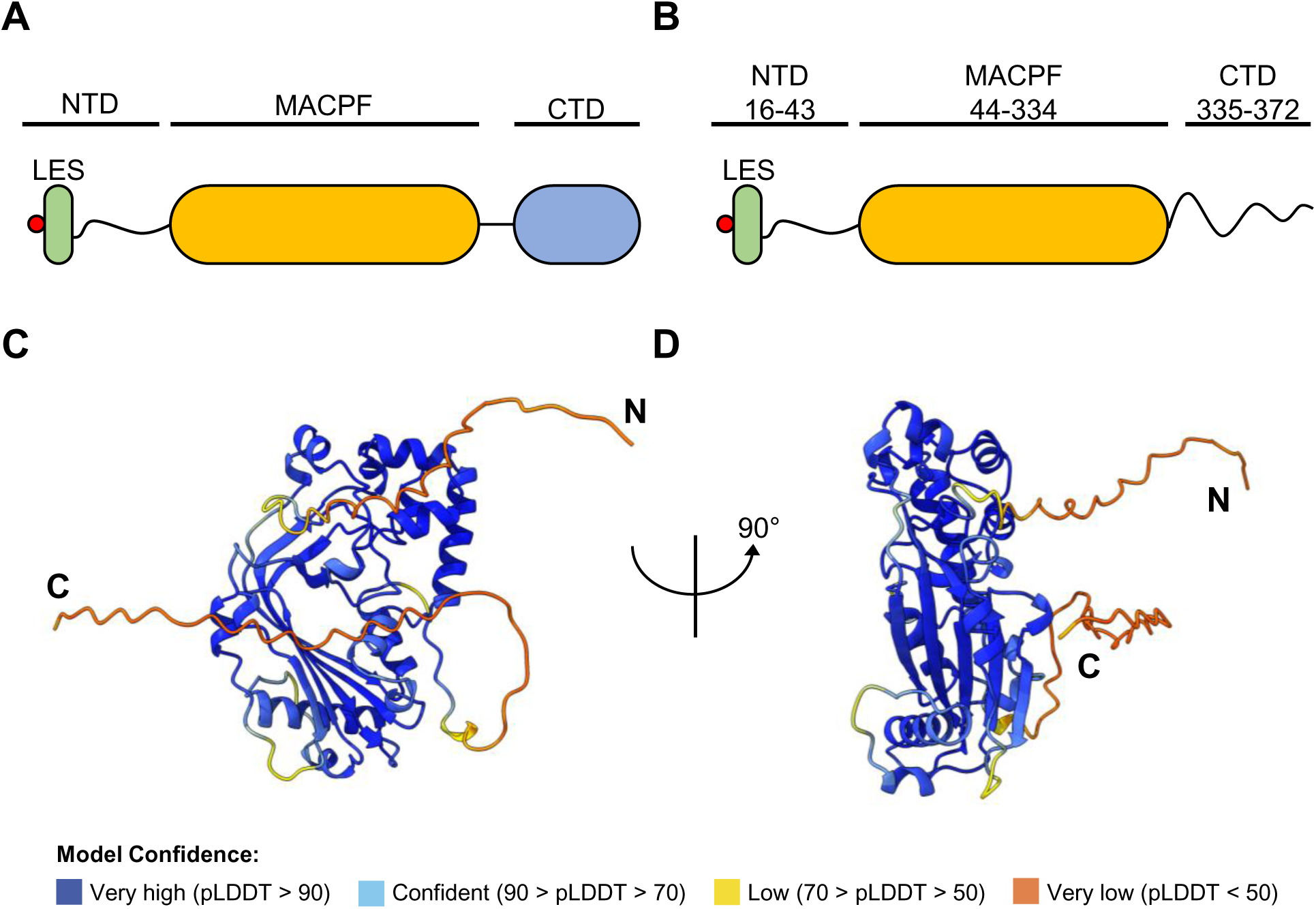
BSAP proteins are structurally organized into three distinct domains. *A*, domain architecture of a generic BSAP (*A*) and of *B. fragilis* 638R BSAP-1 (GenBank: CBW22174.1; *B*). Domain boundaries for BSAP-1 are indicated. The MACPF domain is in *yellow*; the CTD is in *blue*; the LES is in *green*; the *red* circle denotates the lipidated N-terminal cysteine. *C* and *D*, AlphaFold2 model of BSAP-1. pLDDT is AlphaFold’s per-residue confidence score, which scales from 0 to 100.

Here, we explore this hypothesis and show that CTDs are indeed critical for BSAP receptor recognition. To this end, we used BSAP-1 as our model protein, as it has been shown to actively promote strain exclusion in the mammalian gut by targeting the B1R^S^ receptor expressed by sensitive strains that is crucial for mammalian gut colonization (12,13). Using a combination of *in vivo* and biochemical assays, we demonstrate that the CTD of BSAP-1 is both essential and sufficient to promote receptor binding. We expand this finding by performing structural comparison of hundreds of BSAP CTDs, resulting in the clustering of BSAPs according to the structure of their CTD. We additionally uncover that CTD structure can be correlated to their predicted receptor type, reinforcing our finding that CTDs are essential for receptor binding. Lastly, we provide evidence that the BSAP-1 CTD can be repurposed to develop new research tools for the study of Bacteroidales cell biology.

## Results

### The BSAP-1 CTD is essential for *in vivo* bactericidal activity

To determine the importance of CTDs for BSAP functioning, we first tested their relevance for bactericidal activity *in vivo*. To this end, we investigated BSAP-1, produced by *B. fragilis* 638R, which targets the B1R^S^ receptor expressed by other *B. fragilis* strains, including the type strain *B. fragilis* NCTC 9343 (BF9343_1563, GenBank: CAH07344.1). *B. fragilis* 638R itself is protected from BSAP-1 by expressing a homolog of B1R^S^, B1R^R^ (BF638R_1645, GenBank: CBW22173.1), that is not recognized by BSAP-1. AlphaFold2 (33,34) structural modelling of BSAP-1 indicates that the bulk of the mature protein is comprised of the conserved MACPF domain (residues L44-D334) with a short, disordered N-terminal domain (NTD) (residues C16-K43) while, intriguingly, the CTD (residues S335-P372) is only 38 amino acids in length and does not appear to adopt a specific fold (Fig. 1, B, C and D).

To test whether this short polypeptide is indeed sufficient to allow BSAP-1 receptor binding, we chromosomally modified *B. fragilis* 638R to express full-length, C-terminally His-tagged, BSAP-1 (BSAP-1-His) from its native locus as well as a truncated version missing the CTD (BSAP-1-His ΔCTD). Additionally, we engineered a derivative unable to reach the bacterial cell surface by mutating the LES of BSAP-1, substituting the five amino acids immediately downstream of the lipidated cysteine by alanines (residues T17-F21, BSAP-1-His LES*).

To assess bactericidal activity, we took advantage of the fact that the BSAP-1 receptor can be expressed in the heterologous host *Bacteroides thetaiotaomicron*, not targeted by BSAP-1 under normal conditions. We then performed competition assays between *B. fragilis* 638R “predator” strains expressing BSAP-1 or its derivatives and *B. thetaiotaomicron* “prey” strains (*Bt*) expressing the BSAP-1 receptor B1R^S^. Prey survival was assessed by monitoring the growth of serial dilutions on selective media. Importantly, all variants of *B. fragilis* 638R were constructed in a *tssB-tssC* deletion background (hereafter ΔT6SS), leading to inactivation of the T6SS constitutively expressed by this strain (35), allowing us to attribute any observed antagonistic phenotype solely to BSAP expression and function.

Incubation of *B. thetaiotaomicron* expressing B1R^S^ with *B. fragilis* expressing BSAP-1 (wt) or BSAP-1-His led to a ∼3-log reduction in prey survival compared to the untreated control or to cells incubated with a BSAP-1 deletion strain (ΔBSAP-1) (Fig. 2A). Experiments in which *B. thetaiotaomicron* expressed B1R^R^ confirmed that this phenotype is entirely dependent on expression of B1R^S^. As expected, expression of BSAP-1-His LES* led to prey survival similar to that of the untreated control, indicating that surface localization of BSAP-1 is critical for bactericidal activity. Interestingly, expression of BSAP-1-His ΔCTD also resulted in prey survival similar to that of the untreated control, indicating that truncation of the BSAP-1 CTD does prevent bactericidal activity (Fig. 2A). Whole-cell immunoblotting confirmed that the observed phenotypes are likely due to the mutations directly affecting functionality of BSAP-1 rather than expression levels or stability of the protein (Fig. 2B).

**Figure 2.**
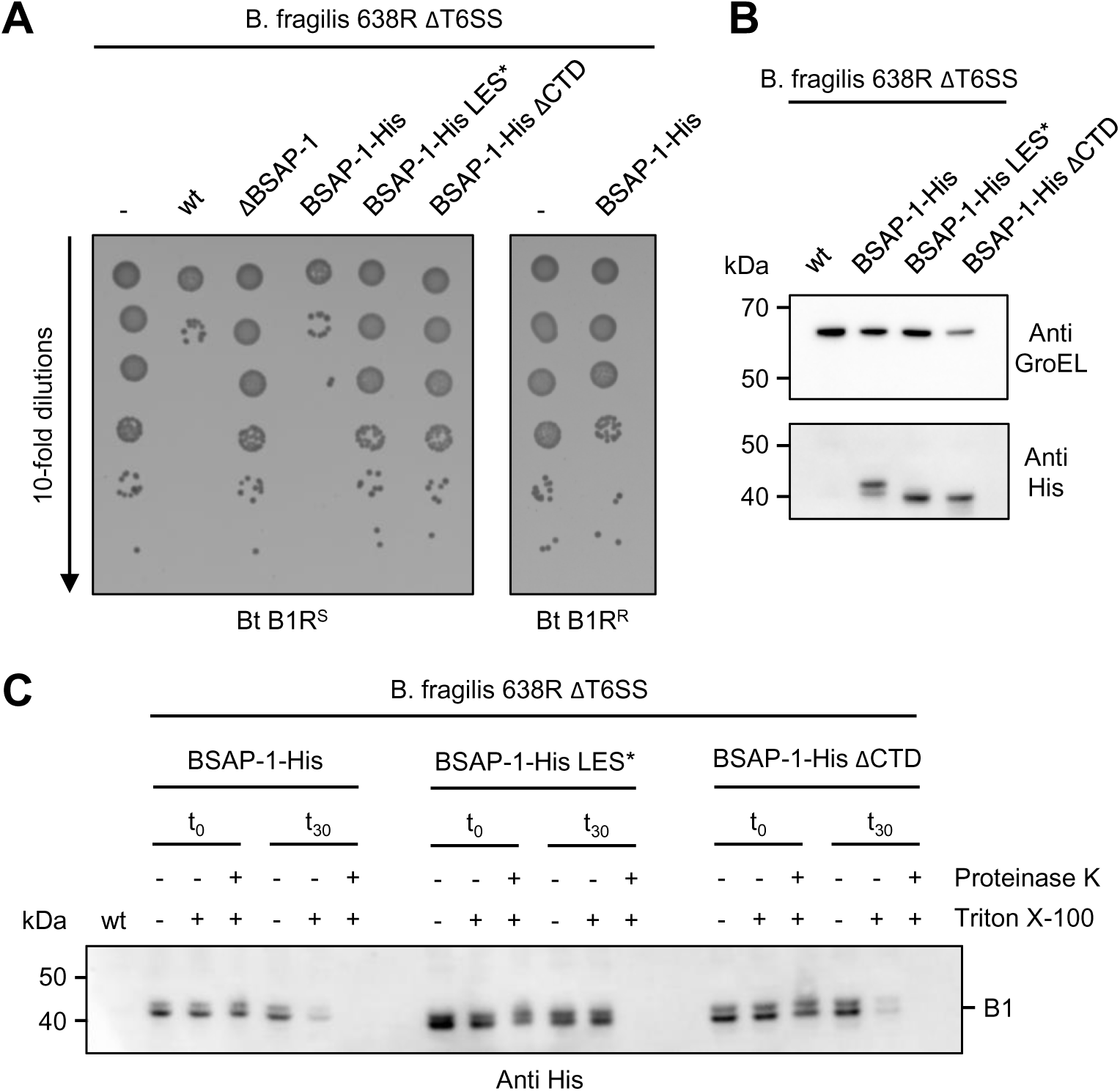
The BSAP-1 CTD is essential for bactericidal activity *in vivo*. *A*, survival of *B. thetaiotaomicron* (*Bt*) prey cells expressing the indicated B1R variant following incubation with various *B. fragilis* 638R predator strains. *B*, whole-cell immunoblot analysis of the indicated *B. fragilis* 638R strains. Detection of GroEL was used as loading control. *C*, immunoblot analysis of the indicated proteinase K-treated *B. fragilis* 638R strains in absence or presence of the detergent Triton X-100. Representative results from at least three independent experiments are shown for each panel.

As evidenced by the LES mutant of BSAP-1, surface localization of the protein is a pre-requisite for bactericidal activity. To exclude the possibility that truncation of the CTD affects proper localization of the protein, we performed Proteinase K surface accessibility assays of cells expressing BSAP-1 and its derivatives. As anticipated, BSAP-1-His was degraded by Proteinase K within 30 min of incubation, both in intact cells and cells permeabilized by the detergent Triton X-100 (Fig. 2C). On the other hand, BSAP-1-His LES* was susceptible to Proteinase K only in the presence of detergent, confirming that the protein is indeed trapped within the cells. BSAP-1-His ΔCTD was readily degraded by Proteinase K in intact cells, demonstrating that deletion of the CTD does not affect translocation of the protein to the cell surface (Fig. 2C). Together, this indicates that the BSAP-1 CTD is not required for correct surface localization of the protein but is critical for bactericidal activity *in vivo*.

### The BSAP-1 CTD is essential for receptor binding

Next, we investigated whether our observed *in vivo* data resulted from the inability of truncated BSAP-1 to bind its receptor or if formation of lytic pores was prevented. To this end, we recombinantly expressed and purified N-terminally His-tagged BSAP-1 as a soluble protein from *E. coli*, omitting the N-terminal signal peptide and lipidated cysteine (data not shown). As we were unable to generate a functional tagged fusion of the BSAP-1 receptor for expression in *E. coli*, we resorted to over-expression of the untagged protein in *B. thetaiotaomicron* followed by cell lysis and recovery of the total membrane fraction containing the receptor. After confirming bactericidal activity of purified BSAP-1 (Fig. S1), we evaluated BSAP-receptor complex formation by performing a pull-down assay using BSAP-1 and incubating it with the B1R^S^-enriched membrane fraction. Following affinity purification, the elution fraction was injected onto a size-exclusion chromatography column and peak fractions were analyzed for presence of BSAP-1 and its receptor. Doing so, we observed two distinct, well-resolved peaks (Peak 1 and Peak 2) (Fig. 3A). SDS-PAGE analysis indicated that Peak 2 contained excess free BSAP-1, while Peak 1 showed that BSAP-1 co-eluted with a protein of approx. 60 kDa, in accordance with the theoretical molecular weight of B1R^S^ (62 kDa) (Fig. 3B). Proteomics analysis confirmed that this band indeed corresponded to B1R^S^ (Table S1), clearly demonstrating the formation of a stable BSAP-1-receptor complex.

**Figure 3.**
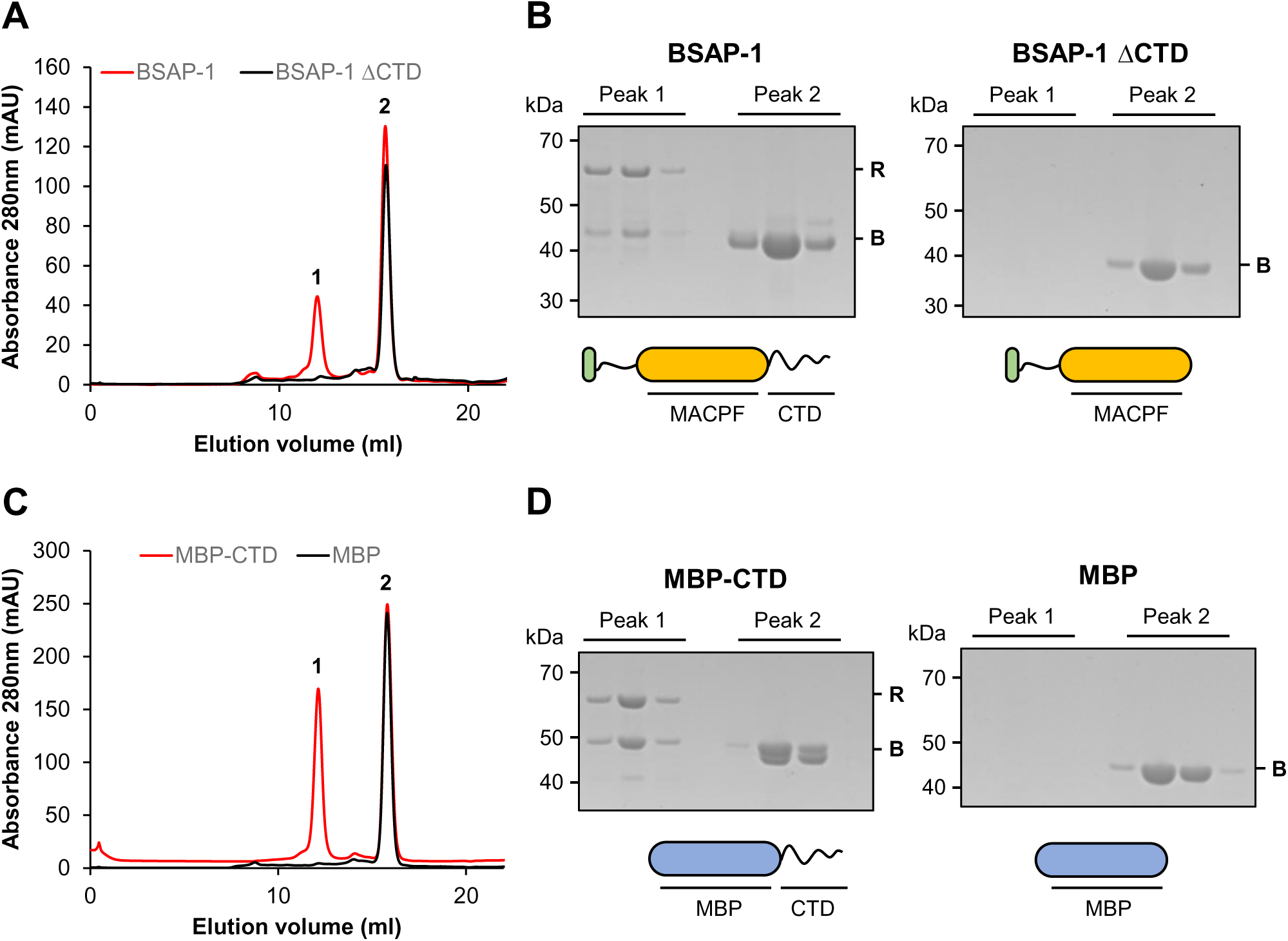
The BSAP-1 CTD is essential and sufficient for receptor binding. *A*, size-exclusion chromatography profile of affinity purified BSAP-1 (*red*) or BSAP-1 ΔCTD (*black*) following incubation with a B1R^S^-containinig membrane fraction. *B*, Coomassie-stained gel showing the protein content of peak 1 and peak 2 from *A* for BSAP-1 (*left*) and BSAP-1 ΔCTD (*right*), respectively. Schematic representation of each protein is indicated below. MACPF domain is in *yellow*; LES is in *green*. *C*, size-exclusion chromatography profile of affinity purified MBP-CTD (*red*) or MBP (*black*) following incubation with a B1R^S^-containinig membrane fraction. *D*, Coomassie-stained gel showing the protein content of peak 1 and peak 2 from *C* for MBP-CTD (*left*) and MBP (*right*), respectively. Schematic representation of each protein is indicated below. MBP is in *blue*. B: bait protein; R: BSAP-1 receptor. Representative results from at least three independent experiments are shown for each panel.

Having successfully validated our experimental approach, we next sought to investigate the effect of CTD truncation on receptor binding. BSAP-1 ΔCTD was purified as above and, unlike purified BSAP-1, did not show any bactericidal activity (Fig. S1). The protein was then used as bait for pull-down assays followed by gel filtration analysis. Only a single peak corresponding to BSAP-1 ΔCTD was observed (Fig. 3, A and B), showing that the protein was unable to bind its receptor and strongly suggesting that the CTD is essential for BSAP-receptor complex formation.

### The BSAP-1 CTD is sufficient for formation of a stable BSAP-1-receptor complex

While the above results supported the idea that the BSAP-1 CTD is directly involved in receptor binding, our results could also be explained by partial misfolding of BSAP-1 following CTD truncation. To rule out this possibility and to clearly demonstrate that the CTD is the sole element required for receptor recognition and binding, we grafted the CTD of BSAP-1 to an unrelated carrier protein, namely *E. coli* Maltose Binding Protein (MBP), resulting in the MBP-CTD fusion protein. Similar to full-length BSAP-1, we observed two distinct peaks (Peak 1 and Peak 2) following pull-down and gel filtration analysis when using purified MBP-CTD as bait in our assay (Fig. 3C). As previously, Peak 2 contained excess free MBP-CTD, while mass spectrometry confirmed that the BSAP-1 receptor B1R^S^ and MBP-CTD co-eluted in Peak 1 (Fig. 3D and Table S1). Pull-down using a purified MBP control protein established that this interaction is specific and dependent on the presence of the BSAP-1 CTD (Fig. 3, C and D). Taken together, our results demonstrate that the BSAP-1 CTD is the driving force providing receptor specificity and is sufficient for interaction with the BSAP-1 receptor.

### BSAPs can be clustered according to the structure of their CTD

BSAPs are able to recognize vastly different receptor types, such as outer membrane proteins or the O-antigen of LPS, with which they must engage in highly specific interactions in order to eliminate their targets with high selectivity. Having demonstrated that the CTD of BSAP-1 is required and sufficient for this purpose, we wanted to confirm that this is a common feature and that the CTD drives receptor specificity across diverse BSAPs. Specifically, we aimed at determining if BSAPs could be classified according to their CTD and if this could inform on the type of receptor that they target. While previous work used full-length sequences and amino acid identity scores to classify BSAPs (14), we reasoned that a structure-based comparison focusing solely on CTDs, less sensitive to sequence variation, might identify shared features among BSAPs targeting similar receptors. To this end, we recovered the full-length sequences of all proteins with a MACPF domain (PF01823) belonging to the Bacteroidota class from the InterPro database (36). Following multiple sequence alignment and clustering, a representative sequence for each cluster was trimmed to encompass only the CTD and its structure predicted using AlphaFold2 (33,34). Lastly, we performed an all-against-all structural comparison using Foldseek (37), allowing us to classify CTDs according to their structural similarity (Table S2). This resulted in the generation of a network of 205 CTDs split into 8 distinct main clusters (Fig. 4A), with the largest one, Cluster 1, comprised of 110 sequences and accounting for 53% of all sequences.

**Figure 4.**
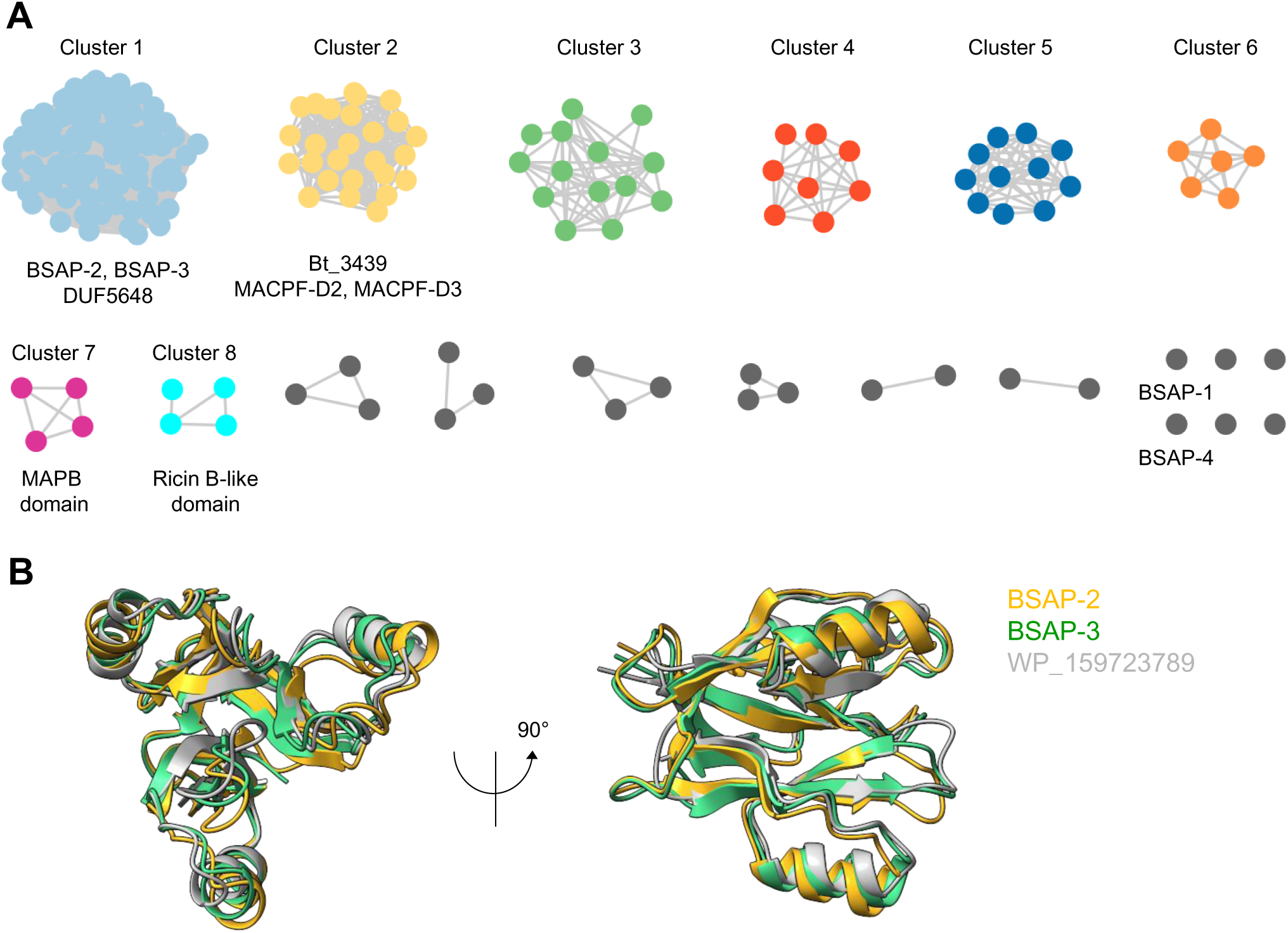
BSAPs can be clustered according to the structure of their CTD. *A*, network analysis of BSAPs clustered according to structural similarity of their CTDs. Previously described BSAPs are indicated below each cluster. Where possible, CTD domain annotation is indicated below each cluster in italicized. *B*, front (*left*) and side (*right*) view of the structural comparison between AlphaFold2 models of the CTD of BSAP-2 and BSAP-3 and the DUF5648 domain of WP_159723789, a previously described GH70 α-glucanotransferase from *Enterococcus* sp. CSURQ0835. BSAP-2 is in *gold*, BSAP-3 is in *green*, WP_159723789 is in *grey*. RMSD with DUF5648: 3,2Å for 129 aligned C^α^ atoms for BSAP-2, 1,6Å for 130 aligned C^α^ atoms for BSAP-3.

Manual inspection revealed that all Cluster 1 CTDs are characterized by a domain of unknown function (DUF5648, Pfam PF18885), widespread throughout the bacterial kingdom and adopting a three-bladed β-α propeller architecture (Fig. 4, B and C). Of note, this cluster contained the CTDs of BSAP-2 and −3, two previously described BSAPs targeting the O-antigen of their respective target strains (13,14), suggesting that other members of this cluster could also target O-antigens. In good agreement with this hypothesis, the DUF5648 domain has recently been identified in a group of α-glucanotransferase enzymes belonging to the wider glycoside hydrolase family 70 (GH70) (Fig. 4, B and C) (38), further indicating that this domain is involved in glycan recognition and binding. This strongly suggests that BSAPs targeting the same type of receptor share an overall similar structure.

The second largest cluster (Cluster 2, 26 sequences) (Fig. 4A) contained the CTD of the only structurally characterized BSAP to date, Bt_3439 (PDB: 3kk7) (32). This CTD is composed of two individual domains of unknown function, D2 and D3 (Pfam PF20779 and PF20785, respectively), that are taxonomically restricted to Bacteroidota. Our Foldseek analysis additionally identified weak structural homologies to MABP for Cluster 7 and to the Ricin B-like lectin domain for Cluster 8 CTDs (Fig. 4A). The MABP domain (MVB12-associated β-prism, InterPro IPR023341) is typically found in eukaryotic proteins in which it acts as membrane-binding module (39). Interestingly, the MABP domain is also found in the Proteobacterial *Photorhabdus luminescens* MACPF protein Plu1415 (PDB: 2QP2) (27). Although non-lytic, Plu1415 was shown to bind to insect cell membranes (27), similar to eukaryotic proteins; BSAPs with this domain could thus target lipids or membranes. The Ricin B-like lectin domain (InterPro IPR035992) is found in numerous proteins throughout the tree of life and is involved in carbohydrate binding (40). Of note, this domain is found in the cytolethal distending toxin subunits A and C produced by several bacterial pathogens, including *E. coli*, *Campylobacter jejuni* and *Helicobacter hepaticus* strains (41,42). The Ricin B-like lectin domain is required for cell membrane binding of subunits A and C and subsequent translocation of cytolethal distending toxin subunit B, resulting in cell death (41,42). As cytolethal distending toxin subunit A is hypothesized to bind glycans, Cluster 7 BSAPs could hence like-wise target carbohydrates. Foldseek analysis of the remaining CTD clusters failed to identify structural similarities to other proteins, suggesting that Bacteroidota have evolved novel protein domains in order to specifically antagonize competitors.

Interestingly, the BSAP-1 CTD appeared as “orphan” and segregated apart from other clusters, likely due to its short, unstructured nature rendering structural comparison difficult. Similarly, the CTD of BSAP-4, also targeting an outer membrane protein (24), failed to fall into a distinct cluster and Foldseek searches did not identify any structural homologs.

### The BSAP-1 CTD can be repurposed to developed new research tools

As we successfully generated an MBP-CTD fusion protein capable of interacting with B1R^S^, we hypothesized that BSAP CTDs could be repurposed for developing new tools for the investigation of Bacteroidales biology. Unlike well-studied organisms such as *E. coli*, little is known about the spatio-temporal organization of the components of the Bacteroidales outer membrane (43). Additionally, the generation of fluorescent derivatives of a protein of interest is often challenging, especially in the case of outer membrane proteins, while metabolic labelling of cell components using click chemistry has so far only been sporadically reported in Bacteroidales (44). As the outer membrane is at the forefront during host-microbe and microbe-microbe interactions, having access to novel high-affinity tools to precisely localize and follow individual membrane components is therefore critical to better understand how these bacteria interact with their environment. As a proof of concept, we therefore grafted the BSAP-1 CTD to HaloTag (HaloTag-CTD), a modified haloalkane dehalogenase designed to covalently bind synthetic ligands, including fluorescent dyes (45). Unlike classical fluorescent proteins like GFP (46), HaloTag does not require oxygen to exhibit strong fluorescence, making it an ideal tool for study of anaerobic bacteria such as Bacteroides. After affinity purification, we first confirmed that the HaloTag-CTD fusion was capable of interacting with B1R^S^ using the above-described pull-down setup (Fig. 5, A and B and Table S1). We also confirmed that this interaction is specific to the CTD by using a HaloTag control protein that did not interact with B1R^S^ (Fig. 5, A and B).

**Figure 5.**
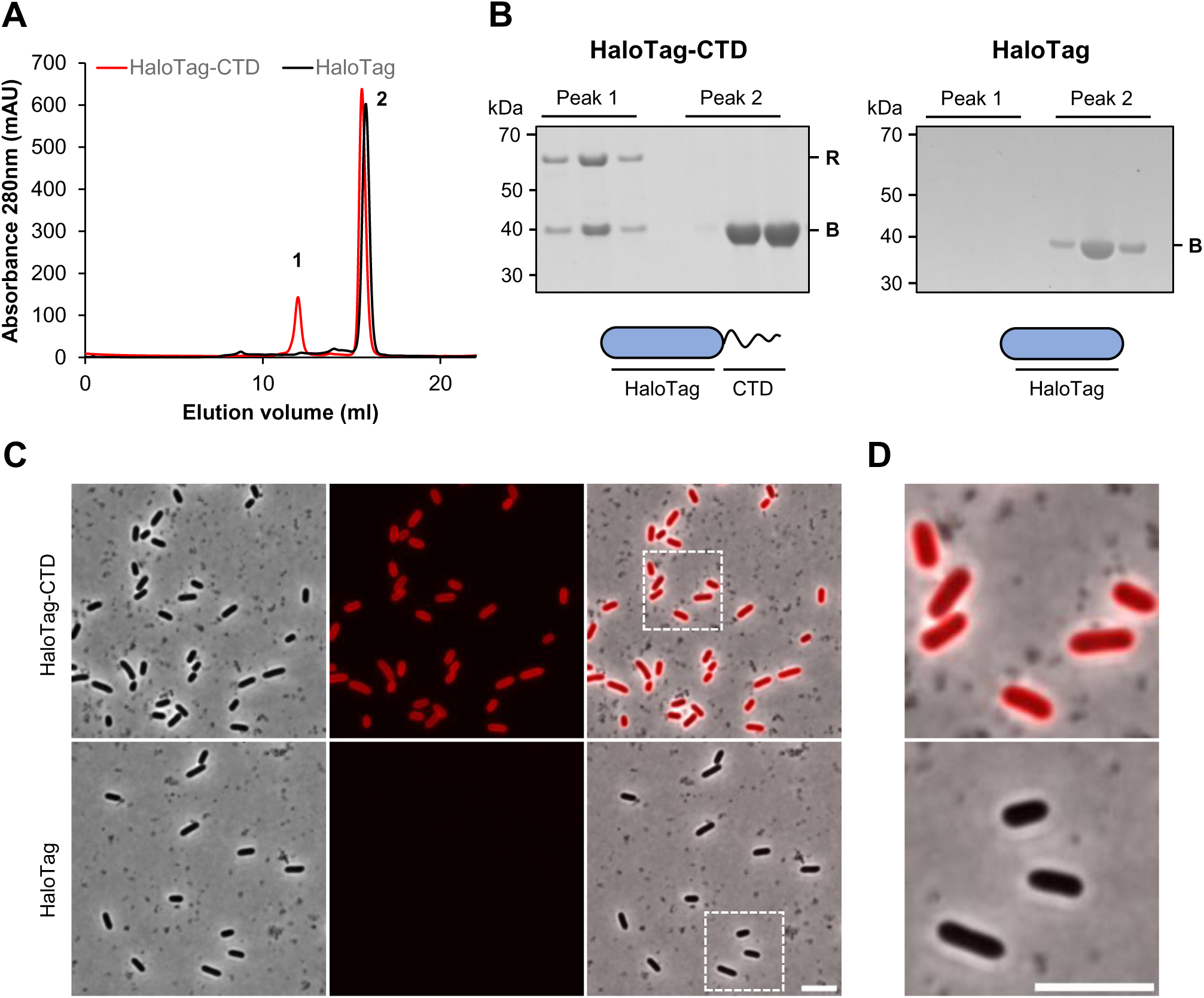
The BSAP-1 CTD can be used to develop new research tools. *A*, size-exclusion chromatography profile of affinity purified HaloTag-CTD (*red*) or HaloTag (*black*) following incubation with a B1R^S^-containinig membrane fraction. *B*, Coomassie-stained gel showing the protein content of peak 1 and peak 2 from *A* for HaloTag-CTD (*left*) and HaloTag (*right*), respectively. Schematic representation of each protein is indicated below. HaloTag is in *blue*. *C*, HaloTag-CTD (*top*) or HaloTag (*bottom*) labelling of *B. thetaiotaomicron* B1R^S^ cells was visualized by epifluorescence microscopy. Scale bar represents 5 µm. *D*, zoom-in of cells boxed in *C*. Scale bar represents 5 µm. Representative results from at least three independent experiments are shown for each panel.

Next, we incubated HaloTag-CTD with exponentially growing *B. thetaiotaomicron* cells expressing B1R^S^, followed by labelling using a HaloTag TMR dye. Fluorescence microscopy inspection of these cells clearly showed strong surface labelling, which was absent from cells labelled with a HaloTag control protein (Fig. 5, C and D) or cells expressing B1R^R^ (Fig. S2, A and B). Identical results were obtained when using *B. fragilis* strain NCTC 9343 endogenously expressing B1R^S^, although with an overall lower fluorescence signal (Fig. S2, C and D). Taken together, these results demonstrate that BSAP-derived probes can be developed and utilized to investigate the outer membrane of Bacteroidales.

## Discussion

BSAPs are a novel class of pore-forming toxins widespread throughout the phylum Bacteroidota that are crucial in mediating bacterial antagonism, including in the human gut. BSAPs are highly specific to their receptor, typically targeting only a single outer membrane protein or LPS variant. This makes them potent weapons for intra-species competition, enabling the producing strain to outcompete its opponents and secure its ecological niche. However, while several BSAP-receptor pairs have been identified, current knowledge falls short in providing insights into how a given BSAP recognizes and binds its receptor with high selectivity.

In this work, we investigated the model protein BSAP-1 and defined which of its domains is involved in providing receptor specificity. We clearly demonstrate that BSAP-1 receptor recognition is entirely driven by the 38 amino acid long CTD of BSAP-1 using a combination of *in vivo* competition assays and *in vitro* protein binding studies. Specifically, we show that deletion of the CTD abrogates BSAP-1 bactericidal activity by preventing receptor binding, while grafting the CTD to unrelated carrier proteins such as MBP and HaloTag enables CTD-driven interaction with the BSAP-1 receptor. Building upon this discovery, we show that BSAPs can be categorized according to the structure of their CTD and, notably, that BSAPs within the same cluster are likely to target the same type of receptor. This study hence lays the groundwork for a more in-depth analysis of BSAPs and how receptor selectivity operates at a molecular level. In this regard, high-resolution structures of BSAP-receptor complexes will represent an invaluable tool to clearly dissect their interactions in the future.

Our data show that the CTD of BSAP-1 can be repurposed to generate tools for the investigation of Bacteroidota biology. Expanding this analysis to other BSAPs, targeting diverse receptors such as the O-antigen of LPS, will provide the opportunity to develop a novel toolbox to study membrane biogenesis in Bacteroidota, a field currently still in its infancy. Indeed, while Bacteroidota are Gram-negative bacteria like the well-studied *E. coli*, several major differences exist regarding outer membrane biogenesis and composition. For instance, Bacteroides species typically produce eukaryotic-like sphingolipids (47), absent from other Gram-negative bacteria. Another notable difference is that unlike most studied bacteria, Bacteroidota display a high percentage of lipoproteins on their surface (26,48). Additionally, Bacteroides species produce penta-acylated, monophosphorylated lipid A, the core component of LPS, in contrast to the hexa-acylated, diphosphorylated lipid A found in *E. coli* (49–51). Lastly, the well-characterized Enterobacterial BAM, Lpt and Lol pathways (52–54) have so far not been investigated in Bacteroidota, raising questions regarding the biogenesis of outer membrane proteins, LPS and lipoproteins, respectively, in this phylum. As a result, virtually nothing is known about how these differences affect outer membrane biogenesis, outer membrane protein composition and spatio-temporal organization, or membrane fluidity. More generally, how Bacteroidota build their cell envelope during cell elongation, whether by zonal, dispersed or polar growth, is currently unknown. Akin to previously described methodologies relying on modified colicins (55), BSAP or BSAP CTD derivatives could hence be used to investigate these many critical aspects of Bacteroidota cell biology, underscoring their potential as research tools.

Our results demonstrate that BSAPs are broadly organized into two distinct domains, with the MACPF domain being responsible for bactericidal activity while the CTD provides receptor specificity. BSAPs therefore exhibit an interesting duality, combining a generic mode of action with high selectivity. Future work focusing on CTD engineering could therefore aim to widen and/or modify the tropism of a given BSAP, enabling it to target and kill various pathogenic species. In light of the global rise of antimicrobial resistance and the resulting threat to human health, having access to molecules specifically designed to target pathogens with high specificity, rather than using broad-range antibiotics, represents an invaluable asset. As BSAPs are able to target both proteins and glycans, two main components of the bacterial outer membrane, BSAPs hence possess a strong potential for the development of novel antimicrobial molecules and warrant further investigation.

## Experimental procedures

### Bacterial strains and growth conditions

All strains and plasmids used in this work are listed in Table S3 and S4, respectively. Bacteroides strains were routinely grown anaerobically in Brain Heart Infusion (BHI, BD) supplemented with 1 g/l cysteine, 5 mg/l hemin and 1 mg/l menadione (BHIs) or modified Gifu Anaerobic Medium (mGAM, Shimadzu Diagnostics Europe) at 37 °C without shaking. *E. coli* strains were routinely grown aerobically in lysogeny broth (LB, MP Biomedicals) at 37 °C with agitation at 200 rpm. Where required, antibiotics were added at the following concentrations: 200 µg/ml ampicillin (Amp) for *E. coli* and 5 µg/ml erythromycin (Em) and 200 µg/ml gentamicin (Gm) for Bacteroides strains.

### Genetic constructs

Plasmids were constructed by standard Gibson cloning or Q5 site-directed mutagenesis (New England Biolabs) using the primers and target DNA listed in Table S5. For heterologous expression of BSAPs in *E. coli*, expression vectors were constructed by cloning the gene of interest, excluding the signal peptide and conserved cysteine, into a pET15b-TEV derivative in frame with a N-terminal 6-His tag and a Ser-Ser-Gly linker. The MPB and HaloTag coding sequences were amplified from pSC95 and *Flavobacterium johnsoniae* FL_147 gDNA, respectively. Bacteroides suicide plasmids to introduce in-frame unmarked deletions were produced by cloning the 500-1000bp upstream and downstream flanking regions of the genes of interest into pSIE1 (a gift from Andrew Goodman, Addgene plasmid #136355; http://n2t.net/addgene:136355; RRID:Addgene_136355) (56). Bacteroides suicide plasmids to introduce chromosomal modifications were constructed by cloning the gene of interest and the 500-1000bp upstream and downstream flanking regions into pSIE1, followed by site-directed mutagenesis. Bacteroides integrative expression plasmids were produced by cloning the gene of interest into pWW3452 in place of sfGFP (57). All plasmid constructs were confirmed by sequencing.

Suicide and expression plasmids were introduced into the appropriate Bacteroides background by biparental mating using *E. coli* S17-1 λpir as donor strain as previously described (58). Briefly, 20 ml mGAM were inoculated at a 1:100 dilution with an overnight culture of Bacteroides and grown anaerobically for 3 h. In parallel, 3 ml LB were inoculated at a 1:100 dilution with an overnight culture of *E. coli* and grown aerobically for 3 h. Cells were collected by centrifugation and washed once with phosphate-buffered saline (PBS). The pellets were then combined, spotted onto BHIs agar and incubated aerobically overnight at 37 °C. The next day, the cells were recovered, resuspended in PBS and serial dilutions plated onto erythromycin and gentamycin containing BHIs agar to select for chromosomal plasmid integration. For in-frame deletion and chromosomal tagging suicide vectors, one of the resulting clones was grown overnight in mGAM without antibiotics to allow for loss of the plasmid backbone, diluted 1:100 into fresh medium, grown for 3-4 h, and then plated on BHIs agar containing 100 µg/l anhydrotetracycline. Anhydrotetracycline-resistant colonies were screened by PCR for the presence of the desired chromosomal modification. All mutant strains were confirmed by sequencing.

### Expression and purification of recombinant proteins

BSAP-1 and its derivatives were purified from *E. coli* BL21 (DE3) cells. The cells were grown in 3.2 l LB medium at 37 °C to an optical density at 600 nm (OD_600_) of 0.5 and protein expression induced by addition of 400 μM IPTG. The cells were then cultured for an additional 3 h at 37 °C. Cells were harvested by centrifugation at 7,500g for 30 min and stored at −20 °C until further use. All purification steps were carried out at 4 °C. Cell pellets were resuspended in buffer A (50 mM HEPES, 150 mM NaCl, 20 mM imidazole, 1 mM DTT, pH 7.5) containing 30 μg/ml DNase I, 400 μg/ml lysozyme and cOmplete™ Protease Inhibitor Cocktail (Merck Life Science) at a ratio of 5 ml of buffer to 1 g of cell pellet. Cells were incubated on ice for 30 min with constant stirring before being lysed by three passages through a French pressure cell at 24,000 PSI. Cell debris was removed by centrifugation at 40,000g for 30 min. The supernatant was then diluted to 125 ml with buffer A, clarified using a Stericup® Quick Release Millipore Express® PLUS 0.22 µm PES filter device (Millipore), and circulated through a 5 ml HisTrap HP column (Cytiva) for 2 h. The column was washed with 12 column volumes (CV) of buffer A and bound proteins were eluted with a 20-500 mM linear gradient of imidazole over 20 CV of buffer A. Peak fractions were collected and concentrated to 2.5 ml using a 10-kDa MWCO Vivaspin® 6 centrifugal filter unit (Sartorius), then injected onto a HiLoad 16/60 Superdex 75 PG column (Cytiva) previously equilibrated in buffer B (50 mM HEPES, 150 mM NaCl, 1 mM DTT, pH 7.5). Peak fractions were concentrated using a 10-kDa MWCO Vivaspin® 6 centrifugal filter unit (Sartorius) and protein concentration was determined using a Microplate BCA™ Protein Assay Kit (Thermo Scientific) according to manufacturer’s instructions. Glycerol was added to a final concentration of 10 % and the proteins were aliquoted into 50 µl fractions before being snap-frozen and stored at −80 °C until further use.

### Isolation of whole membrane fractions

The total membrane fraction of strain sFL153 over-expressing the BSAP-1 receptor was isolated as follows. A pre-culture of sFL153 was grown in mGAM containing erythromycin for 8 h at 37 °C before being used to inoculate 2.4 l of the same medium at a 1:1000 dilution. The cells were then cultured for an additional 16 h at 37 °C. Cells were harvested by centrifugation at 7,500g for 30 min and stored at −20 °C until further use. All subsequent steps were carried out at 4°C. Cell pellets were resuspended in buffer B (50 mM HEPES, 150 mM NaCl, 1 mM DTT, pH 7.5) containing 30 μg/ml DNase I and cOmplete Protease Inhibitor Cocktail (Merck Life Science) at a ratio of 5 ml of buffer to 1 g of cell pellet. Cells were incubated on ice for 30 min with constant stirring before being lysed by three passages through a French pressure cell at 24,000 PSI. Cell debris was removed by centrifugation at 40,000g for 30 min. The supernatant was recovered and total membranes were collected by centrifugation at 180,000g for 60 min. Membranes were resuspended in buffer B and protein concentration determined using a Microplate BCA™ Protein Assay Kit (Thermo Scientific) according to manufacturer’s instructions before being stored at −20 °C until further use.

### BSAP-1 pull-down assay

For BSAP-1 (and derivatives thereof) and BSAP-1 receptor interaction studies, 2 mg of purified protein was mixed with 60 mg of B1R^S^-containing membranes in buffer A in a total volume of 10 ml. The mixture was incubated for 30 min at room temperature with constant agitation. Membranes were then solubilized by addition of n-Dodecyl-β-D-Maltopyranoside (DDM, Anatrace) to a final concentration of 1% (w/v) and incubation for 2 h at 4 °C. Insoluble material was removed by centrifugation at 40,000g for 30 min. The solution was then circulated three times through 500 µl of HisPur™ Ni-NTA Resin (Thermo Scientific). The column was washed with 10 CV of buffer A containing 0.03 % DDM and bound proteins were eluted with 8 CV of Buffer A containing 500 mM imidazole and 0.03 % DDM. The eluate was concentrated to 500 μl using a 10-kDa MWCO Vivaspin® 6 centrifugal filter unit (Sartorius) and then injected onto a Superdex 200 Increase 10/300 GL column (Cytiva) previously equilibrated in buffer B containing 0.03 % DDM. Peak fractions were collected and analyzed by Coomassie staining.

### Competition assays

For competition assays between BSAP-producing predator and BSAP-sensitive prey strains, 5 ml mGAM were inoculated at a 1:100 dilution with overnight cultures of Bacteroides and grown anaerobically for 5 h. One mL of each culture was collected by centrifugation, cells were washed once in PBS, centrifuged again and normalized according to their OD_600_ using PBS. Prey and predators were then mixed at a 1:9 ratio, 5 µl spotted onto BHIS agar, and incubated overnight. The next day, the cells were recovered, resuspended in PBS and serial dilutions spotted onto erythromycin-containing BHIs plates to select for surviving prey cells.

### *In vitro* susceptibility assay

Bactericidal activity of purified proteins was assessed as follows. Five ml mGAM were inoculated at a 1:100 dilution with an overnight culture of Bacteroides and grown anaerobically for 5 h. A total volume of 350 µl of culture was then mixed with 3.5 ml of Top Agar (BHIs supplemented with 7% agar) and poured over a BHIs plate. Sensi-disc™ (BD) impregnated with 15 µg of protein of interest were then deposited on top of the agar, before overnight incubation at 37 °C. The next day, plates were imaged using an ImageQuant LAS 500 Camera (GE Healthcare Life Sciences).

### Proteinase K surface accessibility assay

Surface exposure of BSAP-1 and its derivatives was assessed using a protease accessibility assay. Briefly, 20 ml mGAM were inoculated at a 1:100 dilution with an overnight culture of Bacteroides and grown anaerobically for 5-6 h. Quantities equivalent to 8 ml of OD_600_ = 1 were collected by centrifugation before being resuspended in 640 µl of PBS containing 10 mM MgCl_2_ and 1 mM CaCl_2_.

Cell suspension aliquots of 80 µl were supplemented as appropriate with 200 µg/ml proteinase K (Merck Life Science) and 1 % v/v Triton X-100 (Merck Life Science) and incubated for 30 min at room temperature. Reactions were stopped by the addition of 5 mM phenylmethylsulfonyl fluoride (PMSF, Merck Life Science) and incubation for 5 min, followed by the addition of SDS-PAGE sample buffer containing 1 mM DTT and further incubation at 96 °C for 10 min before analysis by immunoblotting.

### Immunoblotting

The following commercial antisera were used: anti-HisTag peroxidase conjugate (1:4000 dilution, A190-114P, Bethyl) and anti-rabbit IgG peroxidase conjugate (1:5000 dilution NA-934, Cytiva). GroEL antibodies (1:10000 dilution) were raised in rabbits against the purified recombinant GroEL protein (59).

For whole cell immunoblots, Bacteroides strains were cultured in mGAM to OD_600_ = 0.5-0.6, normalized by OD_600_, pelleted by centrifugation and resuspended in SDS-PAGE sample buffer. Samples were then heat-denatured for 10 min at 96 °C before being separated on precast polyacrylamide NuPAGE™ Bis-Tris 10% gels (Life Technologies) using NuPAGE™ MES SDS Running Buffer (Life Technologies). Proteins were transferred onto nitrocellulose membranes using iBlot™ Transfer Stacks in an iBlot™ 2 Gel Transfer Device (Life Technologies). The membranes were then blocked with a 2 % milk powder solution in Tris-buffered saline containing 0.1 % Tween 20 (v/v) for 2 h before being incubated with primary and secondary antibodies as required. Proteins of interest were detected using Immobilon Western HRP Substrate (Merck Life Science) and an ImageQuant LAS 500 Camera (GE Healthcare Life Sciences).

### Epifluorescence microscopy

Fluorescence labelling of Bacteroides strains was carried out as follows. Five ml mGAM were inoculated at a 1:100 dilution with an overnight culture of Bacteroides and grown anaerobically for 5 h. One mL of culture was recovered, cells collected by centrifugation and resuspended in 250 µl mGAM. Twelve µg of HaloTag-CTD or HaloTag control protein were added and the cells incubated anaerobically at 37 °C for 30 min. Cells were then washed three times with PBS before addition of 1 µl of a 50 µM HaloTag tetramethylrhodamine (TMR) ligand (Promega) stock solution and further anaerobic incubation for 20 min in the dark. Cells were washed two times in PBS followed by three washes in mGAM. Finally, 1 µl of cells was spotted onto mGAM low-melt agarose pads previously prepared using 65 µl Gene Frames (Thermo Scientific). Phase contrast and fluorescence microscopy images were acquired on a Ti2-E fully motorized inverted epifluorescence microscope (Nikon) equipped with a CFI Plan Apochromat λ DM 100×1.45/0.13 mm Ph3 oil objective (Nikon), a Sola SEII FISH Illuminator (Lumencor), a Prime BSI camera (Photometrics), a temperature-controlled and light-protected enclosure (Okolab), and a filter cube for mCherry (32 mm, excitation 562/40, dichroic 593, emission 640/75; Nikon). Image acquisition was controlled by the NIS-Ar software (Nikon). All fluorescence images were acquired at 37 °C with 30% power output and 100 ms exposure time.

Images were processed with Fiji, using identical settings for fluorescence contrast and brightness across all regions of interest within the same figure panel unless indicated otherwise.

### Mass spectrometry

Protein bands were visualized by Coomassie Blue staining and in-gel digested with Trypsin (Promega). Peptides were extracted with 0.1% trifluoroacetic acid in 65% acetonitrile and dried in a speedvac.

Peptides were dissolved in solvent A (0.1% trifluoroacetic acid in 2% acetonitrile), directly loaded onto a reversed-phase pre-column (Acclaim PepMap 100, Thermo Scientific) and eluted in backflush mode. Peptide separation was performed using a reversed-phase analytical column (Acclaim PepMap RSLC, 0.075 x 250 mm, Thermo Scientific) with a linear gradient of 4%-27.5% solvent B (0.1% formic acid in 80% acetonitrile) for 40 min, 27.5%-50% solvent B for 20 min, 50%-95% solvent B for 10 min and holding at 95% for the last 10 min at a constant flow rate of 300 nL/min on an Ultimate 3000 RSLC system. The peptides were analyzed by an Orbitrap Exploris240 mass spectrometer (Thermo Fisher Scientific). The peptides were subjected to NSI source followed by tandem mass spectrometry (MS/MS) in Exploris240 coupled online to the nano-LC. Intact peptides were detected in the Orbitrap at a resolution of 60,000. Peptides were selected for MS/MS using HCD setting at 30, ion fragments were detected in the Orbitrap at a resolution of 15,000. A data-dependent procedure that alternated between one MS scan followed by MS/MS scans was applied for 3 seconds for ions above a threshold ion count of 1.0×10^4^ in the MS survey scan with 30.0s dynamic exclusion. MS1 spectra were obtained with an automatic gain control target of 4×10^5^ ions and a maximum injection time set to auto, and MS2 spectra were acquired with an automatic gain control target of 5×10^4^ ions and a maximum injection set to auto. For MS scans, the *m/z* scan range was 350 to 1800. The resulting MS/MS data was processed using Sequest HT search engine within Proteome Discoverer 2.5 SP1 against a *Bacteroides fragilis* NCTC 9343 protein database obtained from Uniprot. Trypsin was specified as cleavage enzyme allowing up to 2 missed cleavages, 4 modifications per peptide and up to 5 charges. Mass error was set to 10 ppm for precursor ions and 0.02 Da for-fragment ions. Oxidation on Met (+15.995 Da), conversion of Gln (−17.027 Da) or Glu (- 18.011 Da) to pyro-Glu at the peptide N-term and acrylamide modification of Cys (+71.037 Da) were considered as variable modifications. False discovery rate (FDR) was assessed using Percolator and thresholds for protein, peptide and modification site were specified at 1%.

### Bioinformatic and structural analysis

All sequences possessing a “MAC/Perforin domain” (Pfam PF01823) belonging to the Bacteroidota class were retrieved from the InterPro database (accessed on November 27th, 2023), resulting in a total of 617 sequences. After a filtering step to keep only the 606 sequences with complete N-termini (starting with a Met), the sequences were aligned using Clustal Omega version 1.2.4 (60). The N-terminal and MACPF domains were removed, and the remaining C-terminal domains were clustered using CD-HIT version 4.8.1 (61) at 95% sequence identity, leading to 270 clusters. For the representative sequence of each cluster, a three-dimensional model was generated using the ColabFold (34) implementation of AlphaFold2 (33). Using these structures, a similarity matrix was built based on the fold similarity computed with FoldSeek (37) followed by clustering with Cytoscape version 3.10.1 (62) using a qtmscore of 0.65 as cut-off value.

## Supporting information

This article contains supporting information.

## Acknowledgments

We thank Gaetan Herinckx for technical help with proteomics analysis. We thank Dr Yoann Santin for assistance in epifluorescence microscopy data acquisition. We thank Dr Francesco Renzi (University of Namur, BE) for sharing strains. We thank Dr Alexandra Gennaris and Dr Laurie Thouvenel for stimulating discussions.

## Author contributions

S.B. and F.L. performed the experimental work unless otherwise indicated. B.I. performed the *in silico* structural analysis. V.D processed the proteomics data. All of the authors discussed the data. S.B., JF.C and F.L. wrote the manuscript. F.L. conceptualized the study. JF.C. and F.L. supervised the study. JF.C. and F.L. secured funding.

## Funding and additional information

This work was funded by the Fonds de la Recherche Scientifique (FNRS) grant agreements WELBIO-CR-2019C-03, the Excellence of Science in Research Program of the FWO and FRS-FNRS (O008818F) and the Fédération Wallonie-Bruxelles (ARC 17/22-087). F. L. was funded by the FRS-FNRS grant 1B.274.22F and by a FWB Incentive Grant.

## Conflict of interest

The authors declare no conflict of interest with the contents of this article.

---

**Figure S1.**
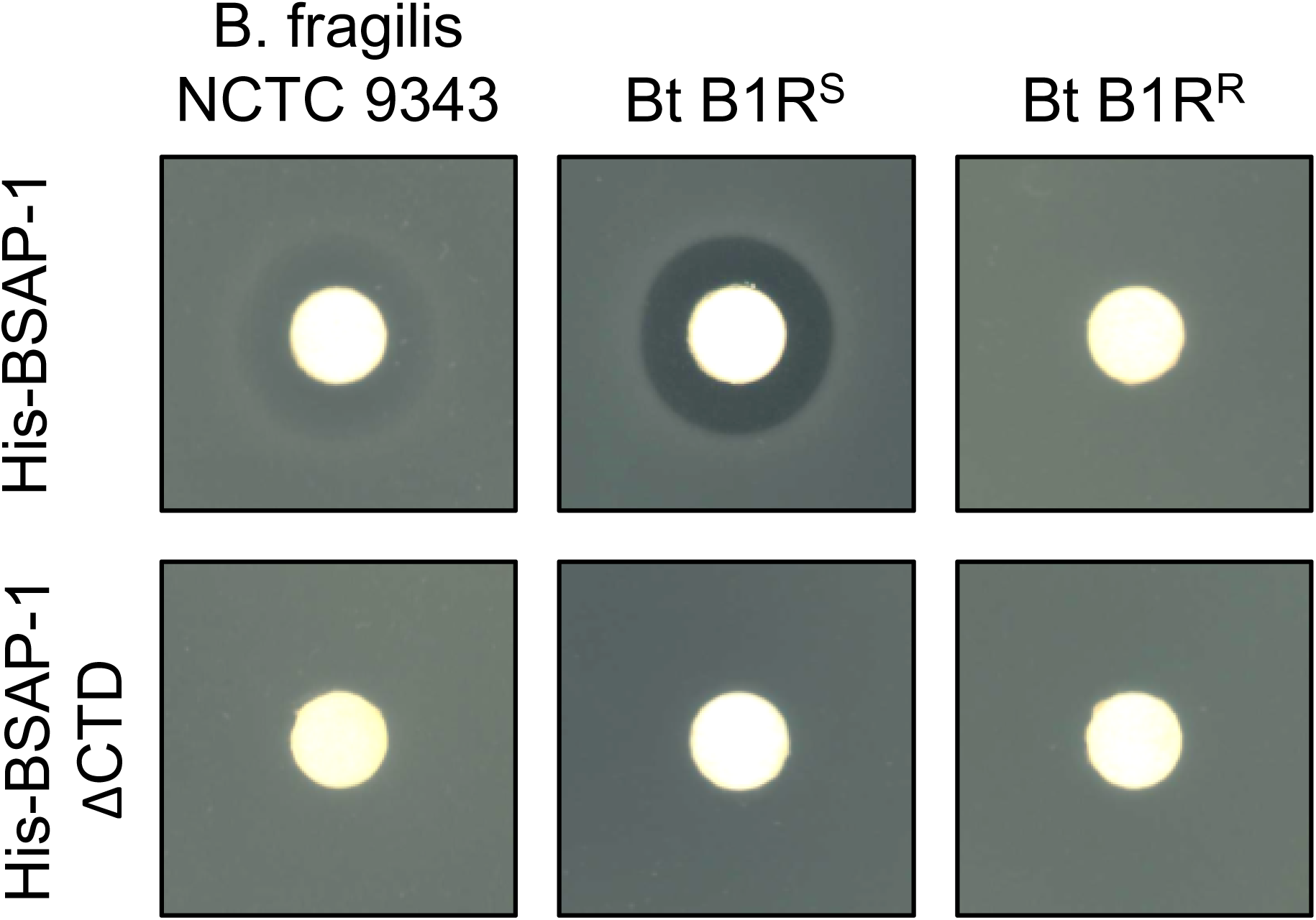
Activity assay of purified recombinant His-BSAP-1 and His-BSAP-1 ΔCTD. Growth inhibition of *B. fragilis NCTC 9343*, *B. thetaiotaomicron* B1R^S^ and *B. thetaiotaomicron* B1R^R^ strains grown in presence of 15 µg of purified His-BSAP-1 (*top row*) or His-BSAP-1 ΔCTD (*bottom row*). Representative results from at least three independent experiments are shown for each panel.

**Figure S2.**
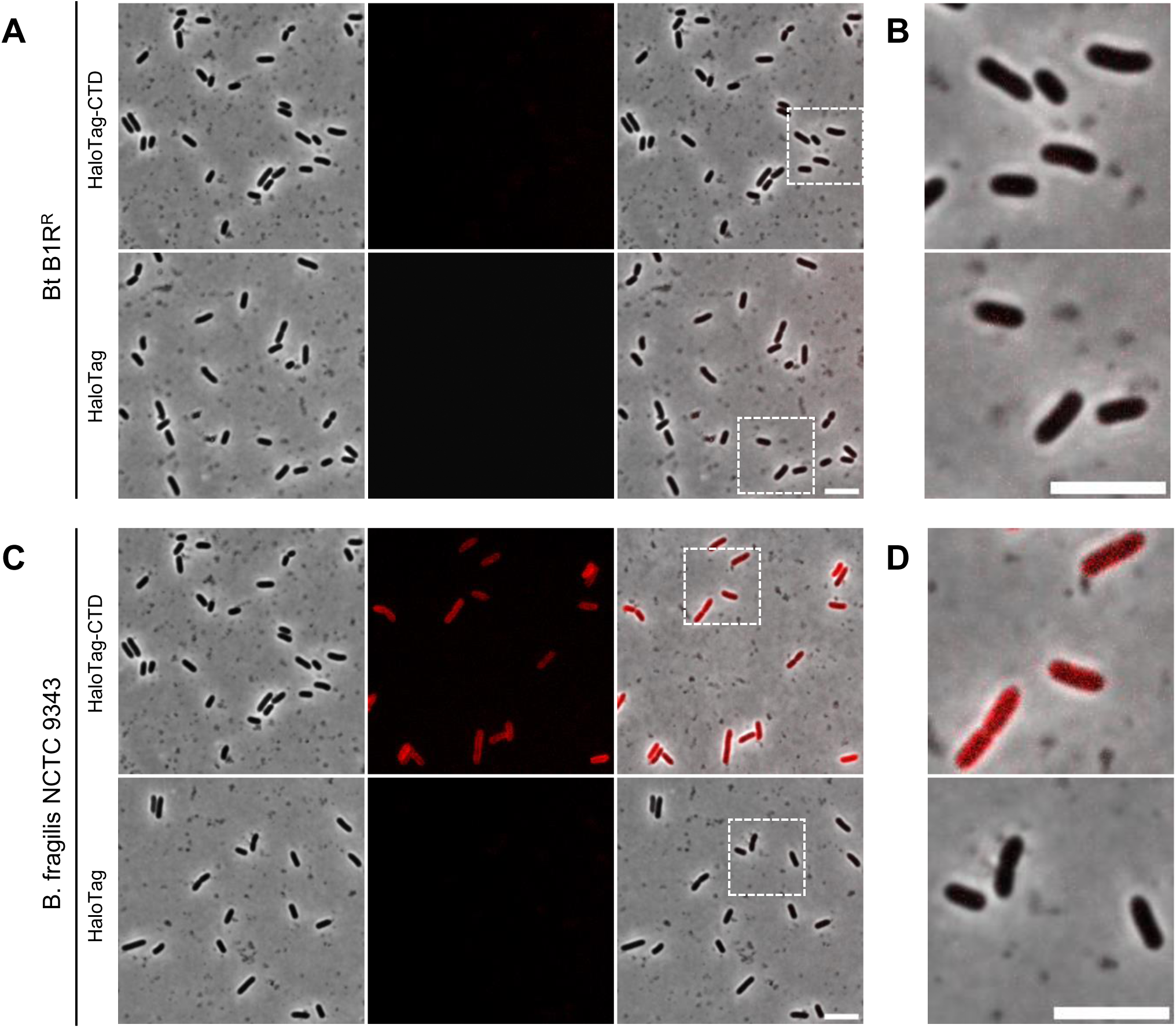
The HaloTag-CTD labelling is specific to cells expressing B1R^S^. *A*, HaloTag-CTD (*top*) or HaloTag (*bottom*) labelling of *B. thetaiotaomicron* B1R^R^ cells was visualized by epifluorescence microscopy. Scale bar represents 5 µm. *B*, zoom-in of cells boxed in *B*. Scale bar represents 5 µm. *C*, HaloTag-CTD (*top*) or HaloTag (*bottom*) labelling of *B. fragilis* NCTC 9343 cells was visualized by epifluorescence microscopy. Scale bar represents 5 µm. *D*, zoom-in of cells boxed in *C*. Scale bar represents 5 µm. Representative results from at least three independent experiments are shown.

## Notes

### Competing Interest Statement

The authors have declared no competing interest.

